# AnDi-J: diffusion analysis of single-particle tracking data in Fiji

**DOI:** 10.64898/2026.10.14.756868

**Authors:** Gorka Muñoz-Gil, Pere Patón-González, Roger Pons-Lanau, Maria F. Garcia-Parajo, Felix Campelo, Carlo Manzo, Francesco Baschieri

## Abstract

Characterizing how particles diffuse is key to understanding biology across scales, yet extracting the underlying physical properties typically requires specialized tools rarely accessible to biology labs. Here we introduce AnDi-J, a Fiji plugin combining classical estimators with state-of-the-art machine learning for single-trajectory analysis and segmentation, requiring minimal knowledge of stochastic processes. It extracts diffusion coefficients and anomalous exponents, classifies trajectories into diffusion models, and segments heterogeneous trajectories into diffusive states, through powerful, easy-to-use visualization tools. See more at https://github.com/gorkamunoz/andi-j.

---

The modeling of biophysical processes as stochastic processes has helped us understand complex systems. In this context, single particle tracking (SPT) [1] has become instrumental for investigating the dynamics of a broad range of biological processes (Fig. 1a), from transcription factor binding to DNA [2] and membrane receptor confinement [3], to diffusion through molecular crowds [4], motor-driven transport [5] and whole-cell migration [6].

**Figure 1.**
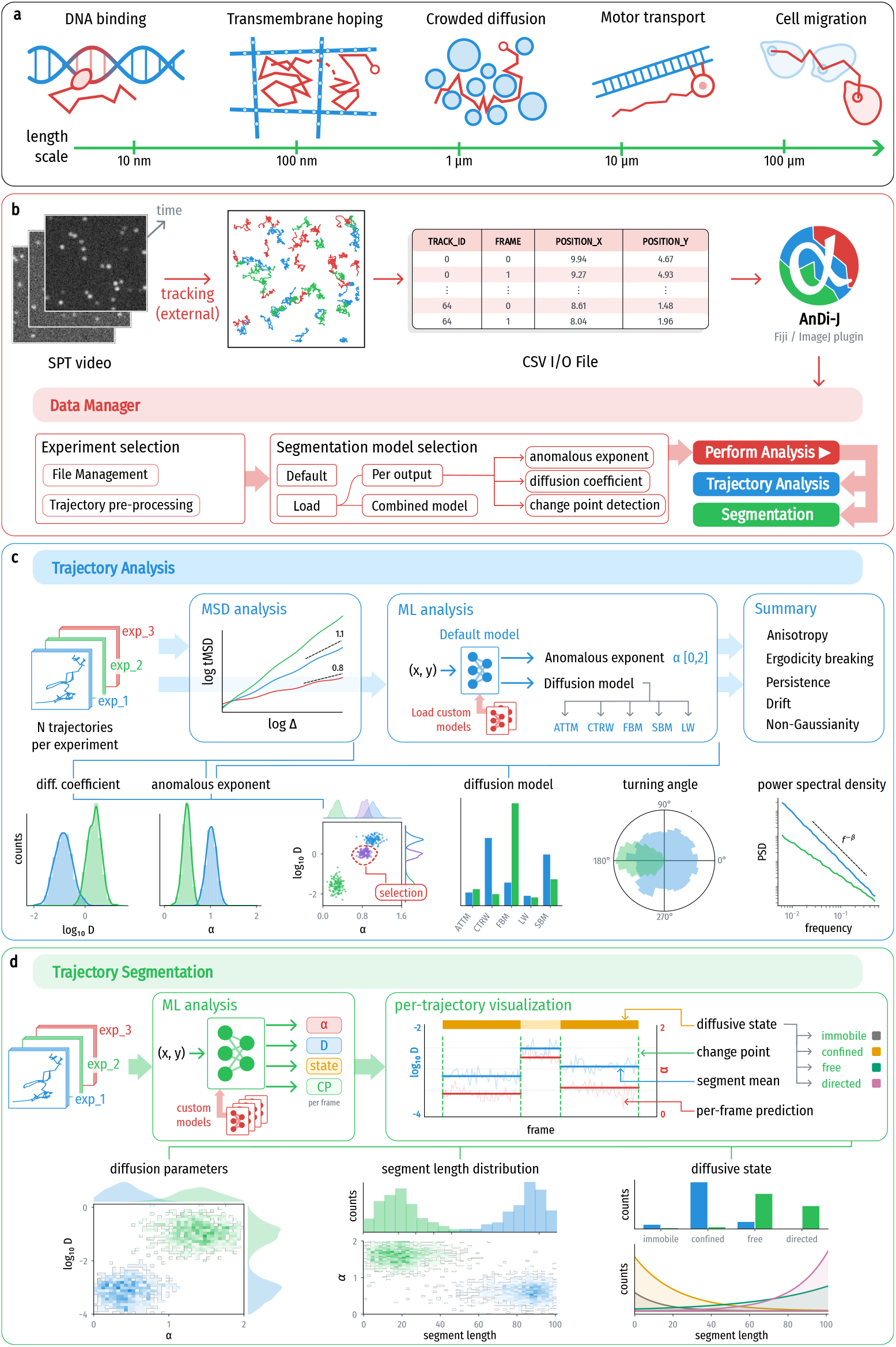
Overview of AnDi-J. **a**, Paradigmatic examples of biological processes where diffusion plays a key role, showcasing the broad scale range over which AnDi-J can be applied. **b**, From an SPT video, particle trajectories are obtained and input to AnDi-J as a CSV file. The Data Manager allows users to select the desired files and pre-process them, after which the analysis is performed. When using the Trajectory Segmentation module, users choose at this stage the ML model used for inference. **c**, Summary of the Trajectory Analysis module. The MSD of the input trajectories can be visualized, and either its fitting or an ML pipeline provides the diffusion coefficient, anomalous exponent and diffusion model classification for each trajectory, together with the turning angle and power spectral density. A summary tab gathers further insights on the dynamics. **d**, Summary of the Trajectory Segmentation module. An ML pipeline predicts diffusive properties for every trajectory’s frames, as well as changepoints that identify segments of distinct diffusive properties. The properties of each segment, together with the distribution of segment lengths, are then provided for further analysis.

These processes are typically analyzed from the trajectories obtained from SPT through classical estimators such as the mean squared displacement (MSD), optimal in clean, theoretical scenarios [7] but highly sensitive to short tracks, localization noise, heterogeneous populations and non-ergodic dynamics [8, 9]. A new generation of estimators, mostly based on machine learning (ML), has largely removed this bottleneck [10–13], with further progress sharpened by the two Anomalous Diffusion (AnDi) Challenges [14, 15], community-wide competitions benchmarking dozens of methods on single-trajectory analysis and trajectory segmentation, respectively.

Yet almost none of this reaches the laboratories generating the data. Most of these approaches require expertise in physics to implement them, raising the barrier for biologists running the experiments. Even the open-sourced AnDi Challenge methods require programming skills, computational resources and, in short, a trained user. The computational biology community, by contrast, has built accessible tools such as ImageJ/Fiji [16], giving access to powerful methods, from image segmentation [17] to particle tracking [18], with minimal prior programming knowledge or formal training in physics. Diffusion analysis tools in Fiji, whether standalone [19, 20] or within wider plugins, typically focus on specific problems or rely on statistical estimators that limit their precision.

AnDi-J fills this gap, bringing state-of-the-art statistical and ML techniques for diffusion analysis to Fiji in a comprehensive yet easy-to-use package. The plugin reads TrackMate-style CSV files [18], integrating seam-lessly into typical biological pipelines. Through the Data Manager window (Fig. 1b), users can load, label and merge files into experiment tags and prepare trajectories for subsequent analysis, providing an organized pipeline for efficiently analyzing large sets of files. The analysis is then divided into two modules, Trajectory Analysis and Trajectory Segmentation, each accompanied by visual tools that let users identify key features in their data and save any prediction or plot for further analysis. All details of the plugin are provided in Ref. [21].

The first module, *Trajectory Analysis*, focuses on single-trajectory analysis (Fig. 1c), starting with the mean squared displacement (MSD) at different levels. Users can visualize the ensemble and time-ensemble average MSD, whose divergence indicates non-ergodic behavior [9], as well as the time-averaged MSD (tMSD) of every trajectory, aiding rapid identification of diffusive populations; *X* and *Y* components can also be plotted separately to detect anisotropic diffusion. Then, linear and logarithmic fits of the tMSD yield the diffusion coefficient *D* and anomalous exponent *α* for every trajectory, with fitting parameters adjustable in case of known localization error or trajectory length [8]. To improve anomalous exponent prediction, known to fail with tMSD fitting on short, noisy or non-ergodic trajectories [9], AnDi-J also offers a state-of-the-art ML pipeline as an alternative which, moreover, also classifies each trajectory into one of five canonical diffusion models (Methods). Through a *D* vs. *α* scatter plot, users can select and save custom subpopulations, treated from then on as a separate experiment in subsequent tabs. The module also provides the turning angle distribution [22], and the power spectral density (PSD), a robust diffusion estimator [23]. Finally, a summary tab gathers the estimated parameters of each experiment in a single exportable table, together with further descriptors of its dynamics: anisotropy, ergodicity breaking, non-Gaussianity, persistence and drift (Methods).

The second module, *Trajectory Segmentation*, uses an ML pipeline to predict, for every frame of every trajectory, the local *α, D* and diffusive state (immobile, confined, free diffusion or directed), together with a changepoint (CP) prediction marking where diffusive changes occur (Fig. 1d). The predicted *α* and *D* are used in the following tabs for further analysis: first, their distribution, letting users identify different diffusive populations in their data; second, the segment-length distribution, combined with these parameters or the predicted diffusive state. This is key to uncovering the dynamical processes underlying particle diffusion, characterizing residence times in specific diffusive states, such as binding or confinement, or capturing continuous changes in diffusive properties over time [24].

To showcase its use on experimental data, we apply AnDi-J to data from three biological systems (Fig. 2a–c). First, we quantify how cells migrate on flat substrates and on 3D nanostructures [25] (Fig. 2a): the tMSD and diffusion coefficient along each axis, together with the turning-angle distribution, show that the nanostructures make migration anisotropic, favoring motion along their direction: the ratio between the diffusion coefficients along the two axes, *D*_*x*_ /*D*_*y*_, given by the summary tab, rises from 1.02 on flat substrates to 2.56 on the nanostructures. Second, we analyze the motion of Halo-KDEL, a small, soluble, engineered protein that localizes in the endoplasmic reticulum (ER) and that transiently visits ER exit sites(ERES), the ER subdomains from where secretory-pathway cargo proteins are exported [26] (Fig. 2b). Trajectory Segmentation identifies mostly trajectories freely diffusing, but also immobilization events. Unlike ERES themselves, mostly immobile or confined at that time scale, the protein diffuses mostly freely, with only a small immobile or confined fraction. Finally, we examine the non-ergodic diffusion of the membrane receptor DC-SIGN [27] (Fig. 2c), evidenced by the divergence between ensemble-averaged and time-averaged MSDs. The estimated frame-wise diffusion coefficients reproduce the wide distribution of *D* characteristic of random diffusivity, with the low-*D* power-law aligning with the annealed transient time motion (ATTM), predominantly predicted by the ML model.

**Figure 2.**
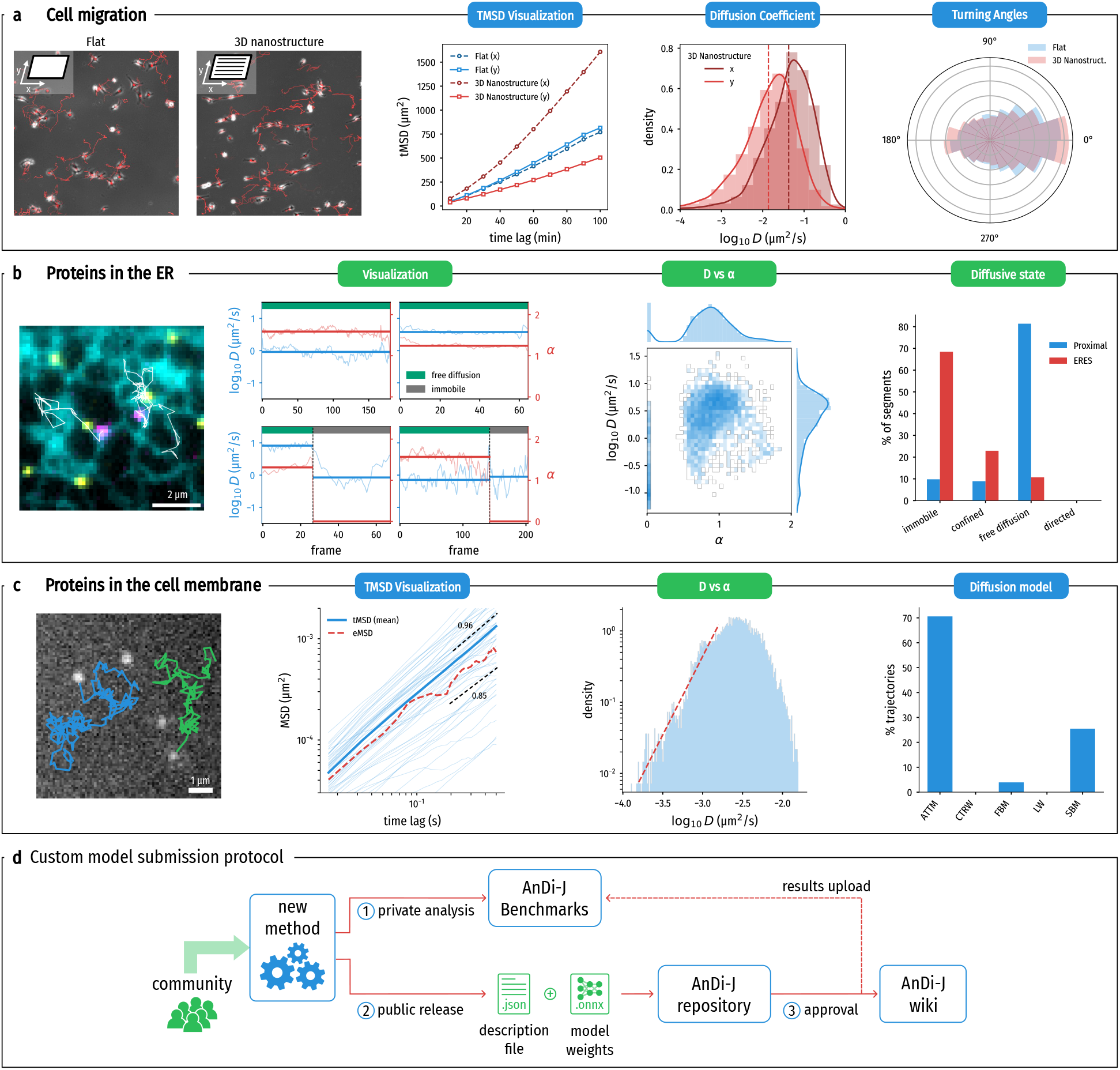
AnDi-J applied to experimental data and community contribution. Blue and green labels mark results from the Trajectory Analysis and Trajectory Segmentation modules, respectively, and name the tab each panel was obtained from. **a**, Cell migration on a flat substrate and on 3D nanostructures aligned with *x* [25]. From left to right: tracked cells, mean tMSD along *x* and *y*, distribution of log_10_ *D* along each axis on the nanostructures (dashed lines: means), and turning-angle distributions. **b**, Halo-KDEL near ERES [26]. From left to right: Halo-KDEL single molecules (magenta, trajectories in white) diffusing along the ER (cyan), and near ER exit sites (yellow); frame-wise (thin) and segment (thick) log_10_ *D* (blue) and *α* (red) for example trajectories, with the predicted diffusive state on top; *D* vs *α* of all frames; and diffusive states of the segments of Halo-KDEL trajectories near ERES (Proximal) and of the ERES themselves. **c**, DC-SIGN receptors in the cell membrane [27]. From left to right: example trajectory; tMSD of every trajectory (cut at 30 time lags), their mean and the eMSD, with log–log fits of the last twenty lags (dashed lines, scaling exponents indicated); distribution of frame-wise log_10_ *D* with a power-law fit at small *D* (dashed line); and ML diffusion model classification. **d**, Community-based development in AnDi-J: (1) developers can privately benchmark a new method on the provided platform (link); (2) they can then release it publicly by open-sourcing their method and submitting a description file through a pull request to the project’s repository; and (3) once approved, the method is documented on the plugin’s wiki and its results are uploaded to the public benchmarks.

AnDi-J is built as an open-source tool bridging two communities: one developing methods for diffusion analysis, the other generating experimental biological data. To this end, AnDi-J promotes the development of new methods that can then be loaded in place of the delivered ones (Fig. 2d). Although many computational methods are public, they typically require expert knowledge, specific software libraries, or have become unusable through version changes. AnDi-J offers researchers a simple way to contribute models that directly benefit the community. The plugin also supports GPU acceleration, automatically leveraging whatever computational resources are available, to speed up inference from ML-based methods.

In conclusion, AnDi-J provides a powerful yet simple tool for experimental biologists to analyze and characterize moving objects, and, as an open, community-driven platform, aims to deepen collaboration between physicists and biologists working on diffusion.

## Methods

We introduce here technical details related to the newly-generated experimental data used in this manuscript, the AnDi-J plugin and the bundled statistical and machine learning (ML) estimators. AnDi-J is an evolving project. The current manuscript is a description of its version v0.1.6, which can be installed in Fiji through the AnDi-J update site (https://sites.imagej.net/AnDi-J/). An up-to-date description of the package is given in its repository (https://github.com/gorkamunoz/andi-j). We also refer the reader to the plugin’s documentation for a step-by-step guide on every tool available (https://github.com/gorkamunoz/andi-j/wiki).

### Cell migration dataset

Polydimethylsiloxane (PDMS - Momentive RTV 615) substrates with or without nanotopographical features were fabricated as previously described in Ref. [25] and manually glued on the bottom of 6-well tissue culture plates. The nanotopographical substrates featured parallel ridges with a height of 0.15 µm and a gap of 1 µm between adjacent ridges. MDA-MB-231 (ATCC HTB-26; a gift from G. Montagnac, Institut Gustave Roussy, Villejuif, France) were cultured in DMEM supplemented with 10% FCS and 100 U/ml penicillin/streptomycin at 37 ^°^C in 5% CO2. Cell cultures were routinely tested for mycoplasma contamination by PCR. For migration experiments, 30,000 cells per well were seeded onto collagen I-coated (50 µg/mL) PDMS substrates. After 24 h, cells were imaged overnight using an epifluorescence microscope (Leica Microsystems Ltd., Wetzlar, Germany) equipped with a 10× Fluotar PH1 objective (NA 0.32) and a PeCon incubation system. Images were acquired at 10-min intervals. When necessary, videos were rotated before tracking analysis to orient the nanotopographical features horizontally. Cell tracking was performed using the TrackMate plugin in Fiji [18].

### Input data and experiment management

Input is one or more CSV files following the standard TrackMate export layout: a header row, three description/unit rows, and one row per spot, with the columns TRACK_ID, POSITION_X, POSITION_Y, FRAME and optionally POSITION_T. Spots are grouped by TRACK_ID and sorted by frame; gaps in FRAME are handled explicitly by all MSD computations. A shared Data Manager loads files either as separate experiments or merged into one, assigns a colour and an editable tag to each, and applies per-file length filters that discard trajectories shorter than a minimum length and truncate longer ones to their first frames up to a maximum length. It reads the spatial calibration and the frame interval of each file from the TrackMate units row, both editable, and the plots and exports can then be shown in physical units. This step is optional and the method falls back in this case to pixels / frames as arbitrary units. Experiments can be added or removed at any point during an analysis session.

### Classical estimators

#### MSD-based estimators

The mean squared displacement (MSD) analysis in the Trajectory Analysis module comprises several levels. We consider an input experiment as a dataset containing *N* 2D trajectories 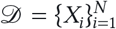, where each trajectory 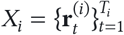 has length *T*_*i*_ and 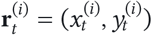. Then, for each trajectory, the time averaged MSD (tMSD) is computed as

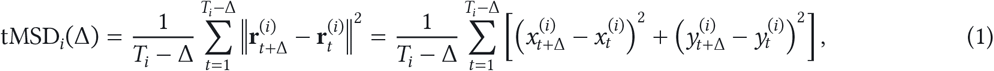

where Δ is the lag time. Given an experiment, its average tMSD is computed as

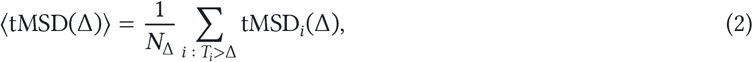

where *N*_Δ_ is the number of trajectories longer than Δ. In addition, the plugin computes the ensemble averaged MSD (eMSD), defined over an experiment as

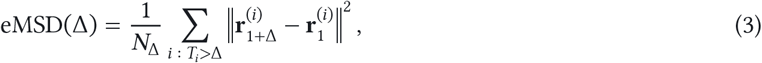

that is, the squared displacement from the initial position after a time Δ, averaged over all trajectories. Comparing the average tMSD and the eMSD is a common measure of the ergodicity of the process [9]. The plugin quantifies it through the ergodicity gap

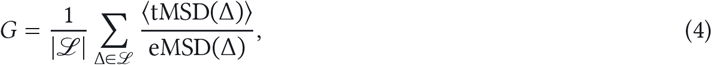

where ℒ is the set of time lags at which both MSDs are computed. *G* ≈ 1 for ergodic processes such as Brownian motion or fractional Brownian motion (FBM), while *G* ≠ 1 indicates weak ergodicity breaking, as in continuous-time random walks (CTRW) or annealed transient time motion (ATTM) [9, 28].

From each of the trajectories’ tMSD, we use a linear and logarithmic fitting to compute the diffusion coefficient *D* and the anomalous exponent *α*, respectively. The user can choose the time lags used for the fitting, allowing users to adapt based on prior knowledge on the localization noise or the trajectory length [8]. The anomalous exponent computed from the logarithmic fitting of the eMSD, preventing any bias arising from non-ergodic diffusion, is also available.

#### Turning angles

Turning angles between successive displacement vectors are computed using

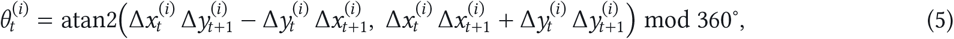

where 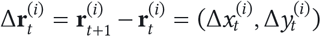, so that 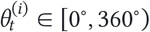, with 0^°^ corresponding to persistent motion, 180^°^ to a full reversal, and angles below (above) 180^°^ to counterclockwise (clockwise) turns. The angles across the whole experiment are then pooled per experiment and displayed as a polar histogram.

#### Power spectral density

The plugin also computes the power spectral density (PSD) of each trajectory as the sum of the one-sided periodograms of its coordinates,

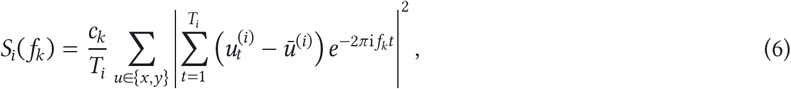

where 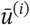 is the mean of coordinate *u* over the trajectory, *f*_*k*_ = *k*/*T*_*i*_ with *k* = 0, …, ⌊*T*_*i*_/2⌋ (in units of inverse frames), and *c*_*k*_ = 2 except for *f*_0_ and, for even *T*_*i*_, the Nyquist frequency, where *c*_*k*_ = 1. The plugin shows, for each experiment, the average of *S*_*i*_ over its trajectories on a log-spaced frequency grid. We refer the reader to Ref. [23] for further details on this method and its interpretation.

## Summary statistics

### Experiment averages

The summary tab gathers, for each experiment, the values computed in the other tabs: the mean *D*, overall and along each axis, the mean *α* obtained from the tMSD fits, from the eMSD fit and from the ML model, the most frequent diffusion model, and the ergodicity gap *G* (Eq. (4)). The ratio *D*_*x*_ /*D*_*y*_ between the mean diffusion coefficients along each axis quantifies anisotropic diffusion, with *D*_*x*_ /*D*_*y*_ = 1 for isotropic motion. The tab also computes the descriptors of the dynamics below, and exports all values as a table.

### Displacement statistics

These descriptors are based on the one-frame displacements 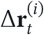, and ⟨⋅⟩ denotes here the average over all displacements of the experiment. The non-Gaussian parameter

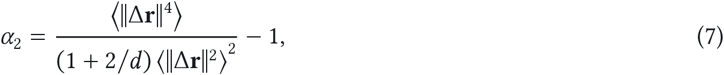

with *d* = 2 dimensions (*d* = 1 when computed along a single axis), vanishes for Gaussian displacements and is positive for heavier tails, which typically arise from heterogeneous diffusion coefficients [29]. The velocity autocorrelation between consecutive displacements,

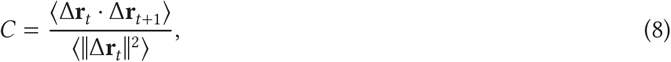

measures persistence: *C* ≈ 0 for uncorrelated steps, as in Brownian motion or CTRW, *C* < 0 for anti-persistent motion, such as FBM with *α* < 1, for which *C* = 2^*α*−1^ − 1, or confinement, and *C* > 0 for persistent, directed or superdiffusive motion [30]. Static localization noise pushes *C* towards −1/2. Finally, the step anisotropy *S* = ⟨cos 2*φ*⟩, with *φ* = atan2(Δ*y*, Δ*x*) the direction of each displacement, measures the alignment of the motion along an axis, with *S* = 1 (*S* = −1) when all steps are along *x* (*y*) and *S* = 0 without a preferred axis.

### Path and drift

These descriptors characterize each trajectory as a whole through its path length 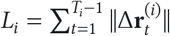 and its net displacement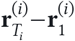, and is then averaged over the trajectories of the experiment [31]. The mean speed is the path length divided by the duration of the trajectory. The forward migration index 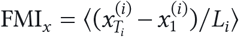, and analogously FMI_*y*_, ranges from −1 to 1 and measures the drift along each axis relative to the path length. The centre of mass 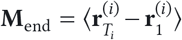 gives the direction and magnitude of the net drift. The directness 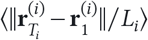 ranges from 0 to 1, the latter for straight trajectories. Since for Brownian motion it decreases as 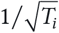, it should only be compared between experiments with similar trajectory lengths. Except for *S* and the directness, all descriptors are also given separately for the *x* and *y* components.

## Machine learning model I/O contracts

We now present the inputs and outputs (I/O) contracts of the different ML models used within the plugin. Any new model loaded by the users (optional) must adhere to this contract and must be in the format of an ONNX file. All models share input size, i.e. a tensor of shape (1, *T*, 2) representing a trajectory of length *T*. The plugin runs the model on several trajectories in parallel. The output depends on the particular variable being predicted. In the Trajectory Analysis module, users can load custom models for either the exponent *α* regression or model classification. In the Trajectory Segmentation module, users can either load a model that predicts all outputs (exponent *α*, diffusion coefficient *D*, diffusive state and changepoints) or only for a specific variable. In the latter case, the rest of the variables fall back to the default model. From the per-frame predictions, the plugin defaults the value of the identified segments to the mean of *α* and *D* and a majority vote on the diffusive state. Custom “all outputs” models can also include their own way of computing the segment level predictions. Details are shown in Table 1. We direct the reader to the plugin’s documentation for further details on the custom model contract.

**Table 1.** Input and output shapes for custom ONNX models. The input is always the raw (*x, y*) positions, one row per frame, with *T* exported as a dynamic axis to allow predictions at any *T*. The *All* segmentation model output is a tuple containing the prediction of the four outputs listed beneath it as separate tensors, each with the shape given in its row. Classes are ordered ATTM, CTRW, FBM, LW, SBM for the diffusion model, and immobile, confined, free diffusion, directed for the diffusive state, the latter given either as probabilities (1, *T*, 4) or as a class index (1, *T*).

| **Tool** | **Variable** | **Input** | **Output** |
|---|---|---|---|
| Trajectory Analysis | Anomalous exponent | (1, *T*, 2) | (1, 1) |
|  | Diffusion model | (1, *T*, 2) | (1, 5) |
| Trajectory Segmentation | All (4 outputs, see caption) | (1, *T*, 2) | — |
|  | *α* | (1, *T*, 2) | (1, *T*) |
|  | log<sub>10</sub> *D* | (1, *T*, 2) | (1, *T*) |
|  | Changepoints | (1, *T*, 2) | (1, *T*) |
|  | Diffusive state | (1, *T*, 2) | (1, *T*, 4) or (1, *T*) |

## Trajectory Analysis default model

Two convolutional networks infer the anomalous exponent and the diffusion model of a trajectory. They share architecture but not weights, and are inspired by the 1st AnDi Challenge team J’s method (based on Refs. [11, 32]). Given an input trajectory, the method first computes its displacements, normalizes them by the mean step size and feeds five parallel blocks, each of four causal convolution layers. The block outputs are concatenated, max-pooled over time and passed to dense layers. Each trajectory runs through two such trunks, as (*δx, δy*) and (*δy, δx*), and the pooled features are merged by an elementwise maximum. Three key improvements have been made to increase the performance of the architecture:

1. The exponent is predicted as a distribution *p*_*i*_ over the 40 values *α*_*i*_ ∈ {0.05, …, 2.00} of the challenge generator, trained against a Gaussian soft target (*σ* = 0.05). It is decoded as the median interpolated within its bin, 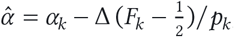, with *F*_*k*_ = ∑_*i*≤*k*_ *p*_*i*_ and Δ = 0.05. The median minimises absolute error, which a scalar output cannot express.
2. Both networks are averaged over the eight symmetries of the axes (sign flips × swap).
3. The exponent posterior *p*_*i*_ enters the classifier as a physical prior. With 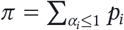, the ATTM and CTRW predicted logits are multiplied by *π* and the Lévy-walk logit by 1 − *π*.

On the 1st AnDi Challenge benchmark for 2D trajectories, compared to the base team J model (shown as FCI in the benchmark page), the exponent MAE fell from 0.152 to 0.137 and the classification *F*_1_ rose from 0.869 to 0.892, making the new method get the first position in both tasks (see Supplementary Figure 1 a).

## Trajectory Segmentation default model

The bundled model is inspired by the winning method of the single trajectory task of the 2nd AnDi Challenge (team I, see [33]). It is based on a one-dimensional U-Net 3+ [34] and predicts *α, D* and diffusive state at each frame, changepoints, and the respective parameters of the resulting segments. We note that the model used within the challenge comprises a set of many parallel models that make it outperform all other submitted methods. However, deploying such a model would be too costly, and we focused on optimizing a single one of these parallel models.

We refined the model in two stages. First, the whole network was retrained without architectural changes, using a reweighted changepoint loss, smoothed changepoint targets and rotation augmentation, and predictions were averaged over eight rotation/reflection views. In the second stage, we froze the main U-Net trunk and focused on improving the heads predicting each of the diffusive parameters:

1. Diffusion coefficient *D*. The original linear regression, which could return negative values, was changed into an MLP that emits log *D*. This makes *D* positive by construction and matches the loss to the log-space metric of the challenge.
2. Anomalous exponent *α*. We adapted the original 1 × 1 readout, which maps the trunk features of each frame to *α* independently of its neighbors, with a residual MLP and a temporal branch of three dilated convolutions that combines features over neighboring frames, since *α* is a long range feature that necessitates displacements across many time lags. The branch also predicts a correction Δ*α* to the per-frame estimate, 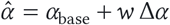 whose optimal weight was found to be *w* = 0.15 using as validation dataset the 2nd AnDi Challenge challenge phase dataset.
3. Diffusive state. The original single layer scalar regression output *s*_*l*_ is now combined with the output of a residual MLP *s*_*m*_ resulting in a single scalar *s* = *s*_*l*_ +*s*_*m*_. Then, we transform this to class probabilities through *p*_*c*_ ∝ exp[−(*s* − *c*)^2^/*T* + *l*_*c*_] (*T* = 0.25), where *l*_*c*_ are per-class logits from a linear readout. This readout was trained on a class-balanced dataset, with additional immobile, trapped and slow subdiffusive trajectories compared to the challenge phase dataset, to ensure that every diffusive state is predicted and the model does not have a bias to predict only free diffusive trajectories. On the 2nd AnDi Challenge benchmark dataset, the F1 score of the immobile state rises from 0 to 0.67, and the mean of the four per-state F1 scores from 0.52 to 0.69.

For segment level prediction, within each segment, *α* and log *D* are aggregated by the median. For the diffusive state, the probabilities *p*_*c*_ are averaged and the most probable state is chosen. Following the labels of the 2nd AnDi Challenge, segments predicted as immobile, and each of their frames, are given *α* = 0, while their *D* is kept as predicted. Compared to the base, original single model approach in Ref. [33], the new method improves in all metrics, and ranks second in the 2nd AnDi Challenge ranking, only behind the multi-network approach of the same authors (see Supplementary Figure 1 b). The final metrics of the provided model, together with its position in the 2nd AnDi challenge metric rank (in parentheses) are: JI (CP) = 0.684 (2), RMSE (CP) = 1.755 (6), MSLE (D) = 0.018 (2), MAE (*α*) = 0.175 (1), F1 (diffusive state) = 0.919 (2).

## Acknowledgments

This work is a direct extension of the AnDi Challenges, and hence we thank all the organizers and participants for their contribution, many of which have been incorporated in the tool with due reference. We also thank Ignacio Arganda Carreras for guidance in the development of the plugin. We acknowledge the extensive use of large language models for the development of the project, from the development of the code underlying the plugin, to writing and editing the different documentation pages and the current manuscript. Every step has been supervised by the authors. G.M.-G. acknowledges funding from the Austrian Science Fund (FWF) [WIT9503323, DOI: 10.55776/WIT9503323], the European Union (ERC Advanced Grant, QuantAI, No. 101055129) and by grant NSF PHY-2309135 to the Kavli Institute for Theoretical Physics (KITP). R.P.-L. and M.F. G.-P acknowledge support from the MICIU/AEI (PID2023-147711NB-I00), Fundació CELLEX (Barcelona), Fundació Privada Mir-Puig, the Generalitat de Catalunya through the CERCA programme. F.C. and R.P.-L. are also supported by “Unidad de Excelencia María de Maeztu” CEX2024-001431-M, funded by MICIU/AEI/10.13039/501100011033. C.M. acknowledges support from grant PID2024-156712NB-I00 funded by MICIU/AEI/10.13039/501100011033 and by “ERDF/EU” and grant AIA2025-164165-C41 funded by MICIU/AEI/10.13039/501100011033. F.B. acknowledges funding from the Austrian Science Fund (FWF) (DOI: 10.55776/PAT4730323).

For open access purposes, the authors have applied a CC BY public copyright license to any author accepted manuscript version arising from this submission. The views and opinions expressed in this article are however those of the author(s) only and do not necessarily reflect those of the European Union or the European Research Council - neither the European Union nor the granting authority can be held responsible for them.

## Code availability

AnDi-J is available at https://github.com/gorkamunoz/andi-j under the MIT License, with documentation at https://github.com/gorkamunoz/andi-j/wiki and the version reported here archived at Ref. [21].

## Data availability

The datasets used to benchmark the AnDi-J default models and new methods through the benchmark webapp are available at Ref. [35].

